# The Ca^2+^ concentration *in vitro* impacts the cytokine production of mouse and human lymphoid cells and the polarization of human macrophages

**DOI:** 10.1101/2021.08.02.454653

**Authors:** Yusuf Cem Eskiocak, Zeynep Ozge Ayyildiz, Sinem Gunalp, Asli Korkmaz, Derya Goksu Helvaci, Yavuz Dogan, Duygu Sag, Gerhard Wingender

## Abstract

Various aspects of the *in vitro* culture conditions can impact the functional response of immune cells. For example, it was shown that a Ca^2+^ concentration of at least 1.5 mM during *in vitro* stimulation is needed for optimal cytokine production by conventional αβ T cells. Here we extend these findings by showing that also unconventional T cells (invariant Natural Killer T cells, mucosal-associated invariant T cells, γδ T cells), as well as B cells, show an increased cytokine response following *in vitro* stimulation in the presence of elevated Ca^2+^ concentrations (approx. 1.8 mM). This effect appeared more pronounced with mouse than with human lymphoid cells and did not influence their survival. A similarly increased cytokine response due to elevated Ca^2+^ levels was observed with primary human monocytes. In contrast, primary human monocyte-derived macrophages, either unpolarized (M0) or polarized into M1 or M2 macrophages, displayed increased cell death in the presence of elevated Ca^2+^ concentrations. Furthermore, elevated Ca^2+^ concentrations promoted phenotypic M1 differentiation by increasing M1 markers on M1 and M2 macrophages and decreasing M2 markers on M2 macrophages. However, the cytokine production of macrophages, again in contrast to the lymphoid cells, was unaltered by the Ca^2+^ concentration. In summary, our data demonstrate that the Ca^2+^ concentration during *in vitro* cultures is an important variable to be considered for functional experiments and that elevated Ca^2+^ levels can boost cytokine production by both mouse and human lymphoid cells.

## Introduction

Various cell media have been developed for *in vitro* cell cultures to optimize the growth and survival of particular cell types. For example, the RPMI1640 media is frequently used for *in vitro* cultures of mouse and human lymphocytes^1–3^. However, it was suggested that the Ca^2+^ concentration of RPMI1640 (0.49 mM) is actually suboptimal for the *in vitro* stimulation of conventional mouse^4^ and human^5^ αβT cells, as measured by cytokine production. Whether the function of unconventional T cells or of other lymphoid and myeloid cells similarly is impacted by the Ca^2+^ concentration *in vitro* is currently unknown. Unconventional T cells differ from conventional αβ T cells by their development and functional capabilities. Prominent examples of unconventional T cells are invariant Natural Killer T (*i*NKT) cells and mucosal-associated invariant T (MAIT) cells, which both express an αβTCR, and γδ T cells, which express a γδTCR. Both *i*NKT and MAIT cells express a highly conserved semi-invariant TCR α-chain, which recognizes glycolipids or riboflavin derivates in the context of the non-polymorphic MHC class I homologs CD1d or MR1, respectively^6,7,8,9^. γδT cells are largely MHC-unrestricted and although the antigen for many γδT cells is not known, some respond to phosphorylated isoprenoid metabolites or lipids^10,11^. These unconventional T cells develop as memory T cells and can provide a first line of defence during immune responses^12^. B cells are the second main adaptive lymphoid cell type and are characterized by the expression of a BCR^13^. As an example of myeloid cells, we choose here macrophages, which are phagocytic and antigen-presenting effector cells of the innate immune system^14^. Depending on the way of stimulation, macrophages can differentiate into several functionally distinct subsets, often referred to as classically activated M1 or alternatively activated M2 macrophages^14,15,16^. To determine the impact of the Ca^2+^ concentration on lymphoid and myeloid cells besides conventional αβT cells, we here compared their immune response *in vitro* in the presence of normal RPMI1640 medium (RPMI^norm^) and RPMI1640 medium supplemented with 1 mM Ca^2+^ (RPMI^suppl^). Our data indicated that elevated Ca^2+^ concentrations during PMA/ionomycin stimulation *in vitro* increased the cytokine production by both mice and human lymphoid cells for most cytokines tested. Furthermore, the polarization of human macrophages shifted towards an M1 phenotype in the high-Ca^2+^ environment. Consequently, the Ca^2+^ concentration during *in vitro* cultures is an important variable to be considered for functional experiments.

## Results

### Cytokine production of mouse unconventional T cells and of B cells is augmented by increased Ca^2+^ concentrations *in vitro*

Following activation, *i*NKT cells are able to produce a wide range of cytokines, including T_h_1 cytokines, like IFNγ and TNF; T_h_2 cytokines, like IL-4 and IL-13, the T_h_17 cytokine IL-17A, as well as IL-10^17,18,19^. When splenic mouse *i*NKT cells were stimulated *in vitro* with PMA and ionomycin, the increased Ca^2+^ concentration had no detrimental effect on *i*NKT cell survival (**Supplementary Figure 1A**). However, we observed a marked and significant increase in the production of all cytokines tested (IFNγ, IL-2, IL-4, IL-10, IL-13, IL-17A) by *i*NKT cells when stimulated in media supplemented with calcium (**Figure 1A-F**). Similar to *i*NKT cells, γδ T cells can produce a wide range of cytokines following stimulation^20^. No detrimental effect of the Ca^2+^ supplementation on γδ T cell survival was observed (**Supplementary Figure 1B)**. However, we noticed a significant increase in the production of IFNγ, IL-2, and IL-4 by γδ T cells stimulated in elevated Ca^2+^ levels (**Figure 2A-C**). In contrast, the changes for IL-10 and IL-17A remained non-significant (**Figure 2D, 2E**). We also analysed the impact of Ca^2+^ levels on B cells. The survival of stimulated B cells was not impaired by the Ca^2+^ concentration *in vitro* (**Supplementary Figure 1C**). However, an increase in the production of IL-2 and IL-10 was observed (**Figure 3B, 3C**), while IFNγ (**Figure 3A**) remained unaffected. Therefore, the PMA/ionomycin stimulation of mouse lymphoid cells in complete RPMI medium supplemented with 1.0 mM Ca^2+^ improves the detection of numerous cytokines, without impacting the survival of the cells.

**Figure 1.**
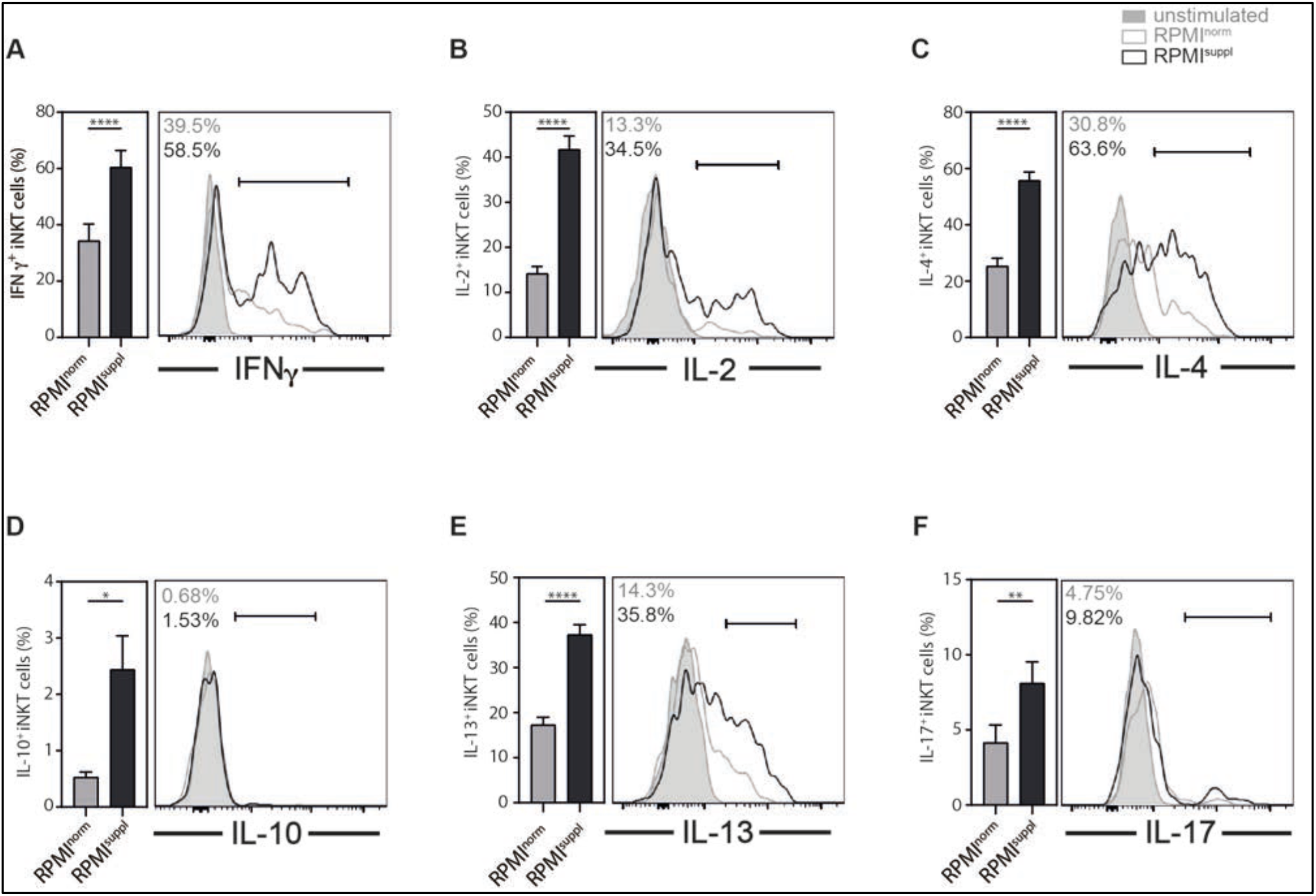
Ca^2+^ supplementation *in vitro* increases the cytokines production of mouse *i*NKT cells. **(A-F)** Splenocytes from C57BL/6 mice were stimulated 4 h with 50 ng/ml PMA and 1 μg/ml ionomycin in either normal RPMI1640 medium (RPMI^norm^) or RPMI1640 medium supplemented with 1 mM Ca^2+^ (RPMI^suppl^). The production of **(A)** IFNγ, **(B)** IL-2, **(C)** IL-4, **(D)** IL-10, **(E)** IL-13, and **(F)** IL-17A by *i*NKT cells (live CD8α^-^CD19/CD45R^-^ CD44^+^ TCRβ/CD3ε^+^ CD1d/PBS57-tetramer^+^ cells) was analysed by intracellular cytokine staining (ICCS). Summary graphs (left panels) and representative data (right panels) from gated *i*NKT cells are shown, respectively. Data were pooled from three independent experiments with three mice per group per experiment (n = 9).

**Figure 2.**
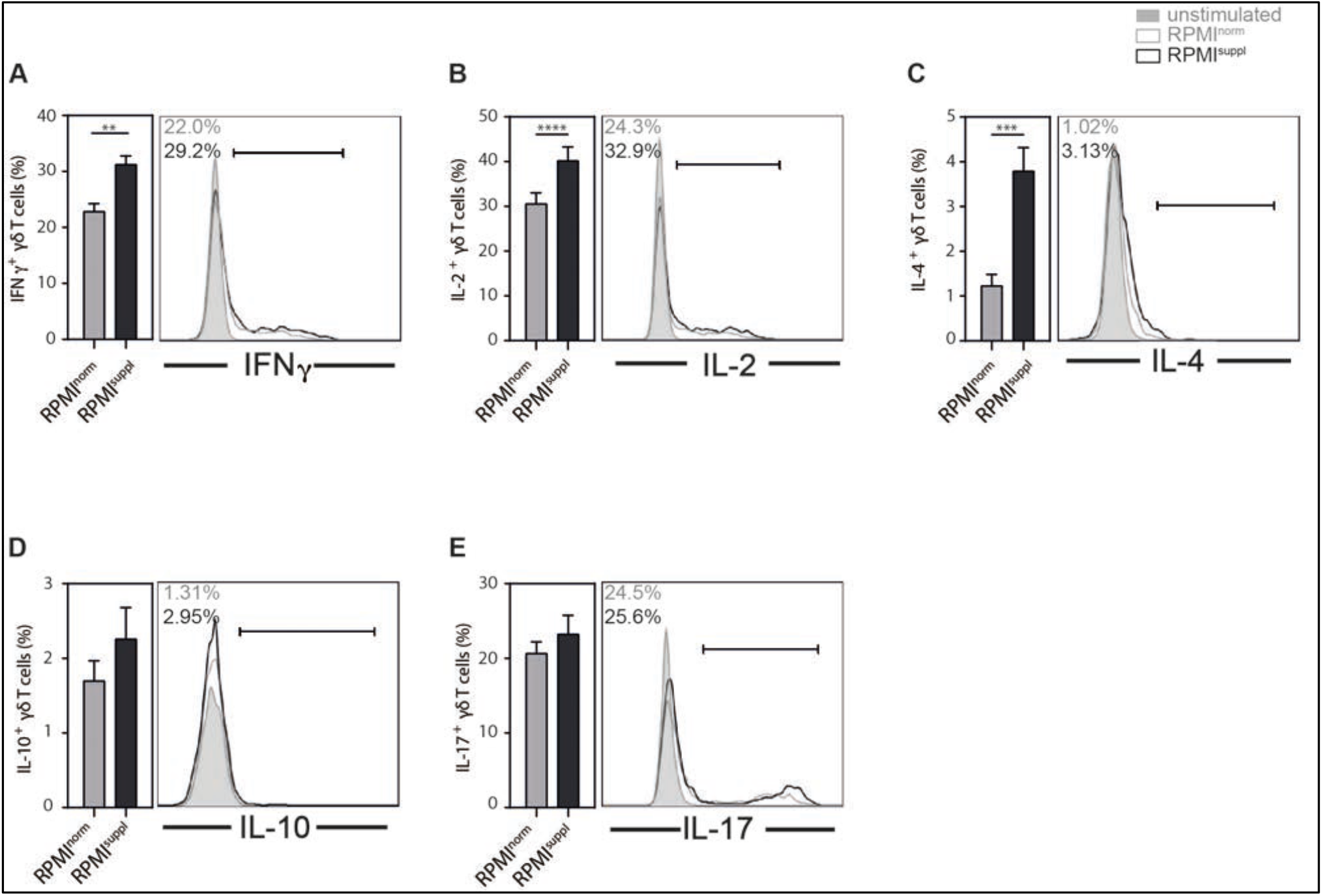
Ca^2+^ supplementation *in vitro* increases the production of some cytokines by mouse γδ T cells. **(A-E)** Splenocytes from C57BL/6 mice were stimulated 4 h with 50 ng/ml PMA and 1 μg/ml ionomycin in either normal RPMI1640 medium (RPMI^norm^) or RPMI1640 medium supplemented with 1 mM Ca^2+^ (RPMI^suppl^). The production of **(A)** IFNγ, **(B)** IL-2, **(C)** IL-4, **(D)** IL-10, and **(E)** IL-17A by γδ T cells (live CD19/CD45R^-^ CD4^-^ CD8α^-^CD3ε^+^ γδTCR^+^ cells) was analysed by ICCS. Summary graphs (left panels) and representative data (right panels) from gated γδ T cells are shown, respectively. Data were pooled from three independent experiments with three mice per group per experiment (n = 9).

**Figure 3:**
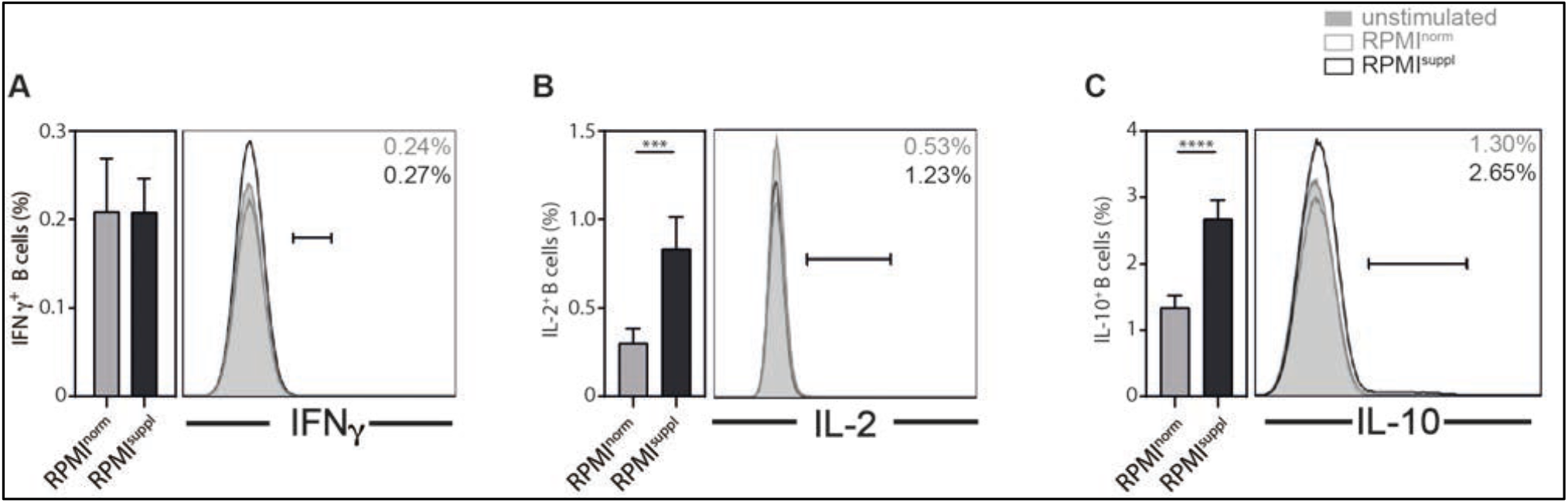
Ca^2+^ supplementation *in vitro* increases the production of IL-2 and IL-10 by mouse B cells. **(A-C)** Splenocytes from C57BL/6 mice were stimulated 4 h with 50 ng/ml PMA and 1 μg/ml ionomycin in either normal RPMI1640 medium (RPMI^norm^) or RPMI1640 medium supplemented with 1 mM Ca^2+^ (RPMI^suppl^). The production of **(A)** IFNγ, **(B)** IL-2, **(C)** IL-10 by B cells (live CD3ε^-^CD4^-^ CD8α^-^CD19/CD45R^+^ cells) was analysed by ICCS. Production of IL-4, IL-13, or IL-17A was not detected (data not shown). Summary graphs (left panels) and representative data (right panels) from gated B cells are shown, respectively. Data were pooled from three independent experiments, with three mice per group per experiment (n = 9).

### Cytokine production of human unconventional T cells and of B cells is augmented by increased Ca^2+^ concentrations *in vitro*

Having established that increased Ca^2+^ concentrations during *in vitro* stimulation can augment the cytokine production of mouse lymphoid cells, we next tested its effect on human lymphoid cells. For primary human *i*NKT cells, the elevated Ca^2+^ levels only increased the production of TNF slightly, without effects on the other cytokines tested (IFNγ, IL-2, IL-4, IL-17A) or on the expression of the activation marker CD69 (**Figure 4A-F**). For primary human Vδ2^+^ T cells, some (IFNγ, TNF) but not all (IL-2, IL-4) were boosted by the Ca^2+^ supplementation (**Figure 5A-E**). For primary human MAIT cells, the elevated Ca^2+^ levels increased the production of all cytokines tested (IFNγ, IL-2, IL-17A), without changes to the expression of CD69 (**Figure 6A-D**). A similar effect of the Ca^2+^ concentrations was noted for primary B cells (IL-2, TNF, CD69) (**Figure 7A-C**). For some immune cells, the low frequency in the peripheral blood makes their analysis directly *ex vivo* difficult, which is why protocols were established to expand them *in vitro*. We, therefore, also tested the impact of elevated Ca^2+^ concentrations on *in vitro* expanded *i*NKT and Vγ2^+^ T cells. For expanded human *i*NKT cells, the elevated Ca^2+^ levels only increased the production of TNF slightly, without effects on the other cytokines tested (IFNγ, IL-2, IL-4, IL-17A) and decreased CD69 expression (**Figure 8A-F**). For expanded human Vγ2^+^ T cells, some (GM-CSF, IFNγ, TNF) but not all (IL-4, CD69) markers were boosted by the Ca^2+^ supplementation, whereas the production of IL-2 surprisingly decreased (**Figure 9A-F**). For all human lymphoid cell populations tested, the Ca^2+^ supplementation *in vitro* did not impair the cell survival (**Supplementary Figure 2A-D**). Therefore, similar to mouse lymphoid cells, the detection of cytokines in primary human lymphoid cells can be improved by increasing the Ca^2+^ levels during the *in vitro* stimulation.

**Figure 4:**
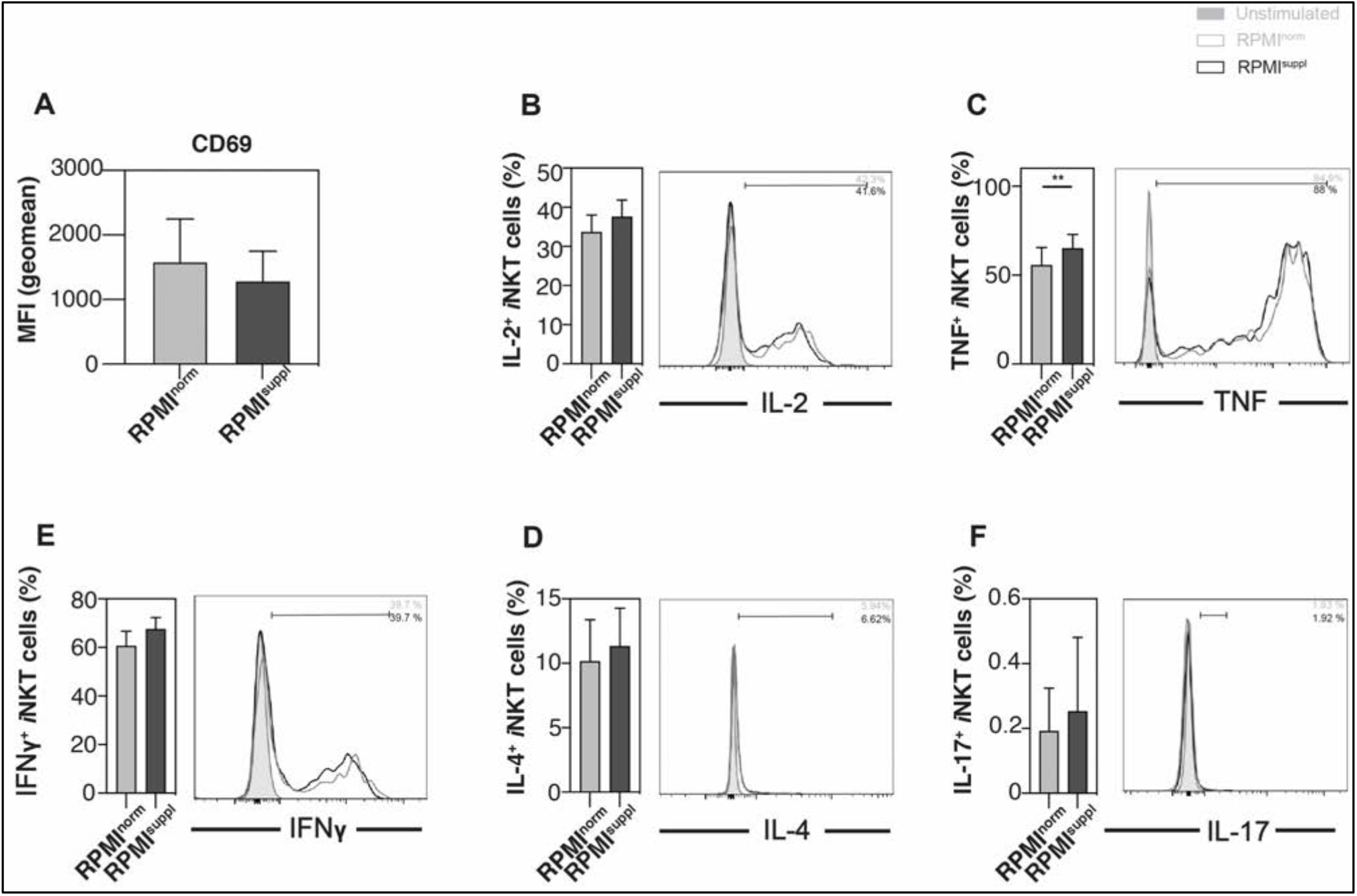
Ca^2+^ supplementation *in vitro* increases TNF production by primary human *i*NKT cells. PBMCs were isolated from the residual leukocyte units of healthy donors. PBMCs were stimulated for 4 h with 25 ng/ml PMA and 1 μg/ml ionomycin in either normal RPMI1640 medium (RPMI^norm^) or RPMI1640 medium supplemented with 1 mM Ca^2+^ (RPMI^suppl^). Human Vα24*i* NKT cells (live CD14^-^ CD20^-^ CD3^+^ 6B11^+^ cells) were analysed for the expression of the activation marker **(A)** CD69 and the production of the cytokines **(B)** IL-2, **(C)** TNF, **(D)** IFNγ, **(E)** IL-4, and **(F)** IL-17. Summary graphs (left panels) and representative data (right panels) from gated *i*NKT cells are shown, respectively. Data were pooled from three independent experiments with three samples each (n = 9).

**Figure 5:**
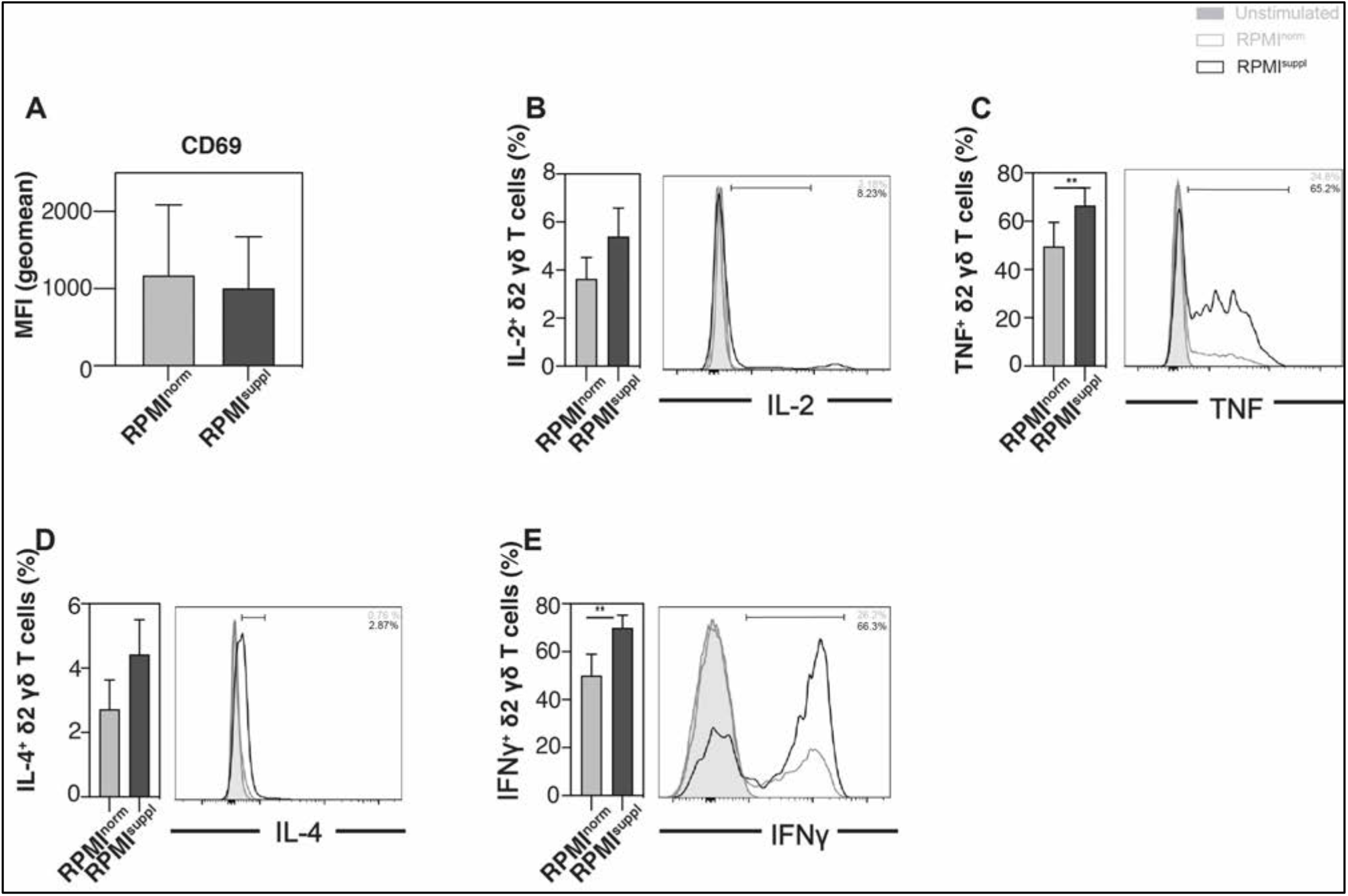
Ca^2+^ supplementation *in vitro* increases the production of TNF and IFNγ from primary human Vδ2^+^ T cells. PBMCs were isolated from residual leukocyte units of healthy donors and were stimulated for 4 h with 25 ng/ml PMA and 1 μg/ml ionomycin in either normal RPMI1640 medium (RPMI^norm^) or RPMI1640 medium supplemented with 1 mM Ca^2+^ (RPMI^suppl^). Human Vδ2^+^ T cells (live CD14^-^ CD20^-^ CD3^+^ γδTCR^low^ or Vγ2^+^ cells) were analysed for the expression of **(A)** CD69 and the production of **(B)** IL-2, **(C)** TNF, **(D)** IL-4, and **(E)** IFNγ. Summary graphs (left panels) and representative data (right panels) from gated Vδ2^+^ T cells are shown, respectively. Data were pooled from three independent experiments with three samples each (n = 9).

**Figure 6:**
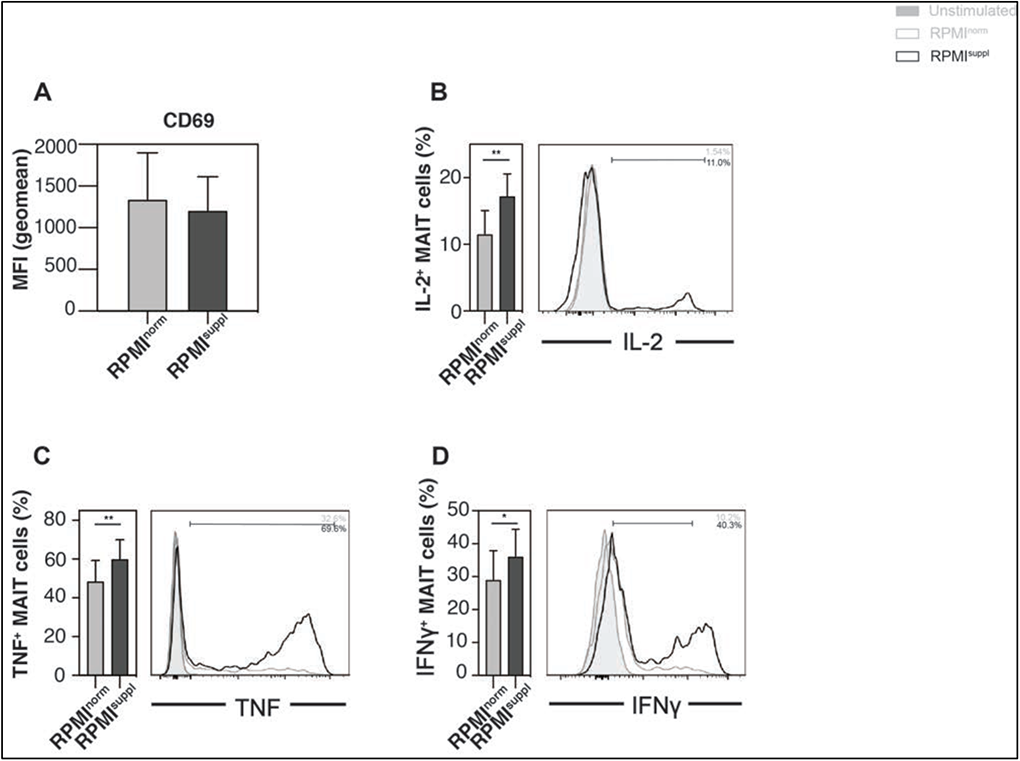
Ca^2+^ supplementation *in vitro* increases the cytokine production of primary human MAIT cells. PBMCs were isolated from residual leukocyte units of healthy donors. PBMCs were stimulated for 4 h with 25 ng/ml PMA and 1 μg/ml ionomycin in either normal RPMI1640 medium (RPMI^norm^) or RPMI1640 medium supplemented with 1 mM Ca^2+^ (RPMI^suppl^). Human MAIT cells (live CD14^-^ CD20^-^ CD3^+^ Vα7.2^+^ CD161^+^ cells) were analysed for the expression of **(A)** CD69 and the production of **(B)** IL-2, **(C)** TNF, and **(D)** IFNγ. Summary graphs (left panels) and representative data (right panels) from gated MAIT cells are shown, respectively. Data were pooled from four independent experiments with three samples each (n = 12).

**Figure 7:**
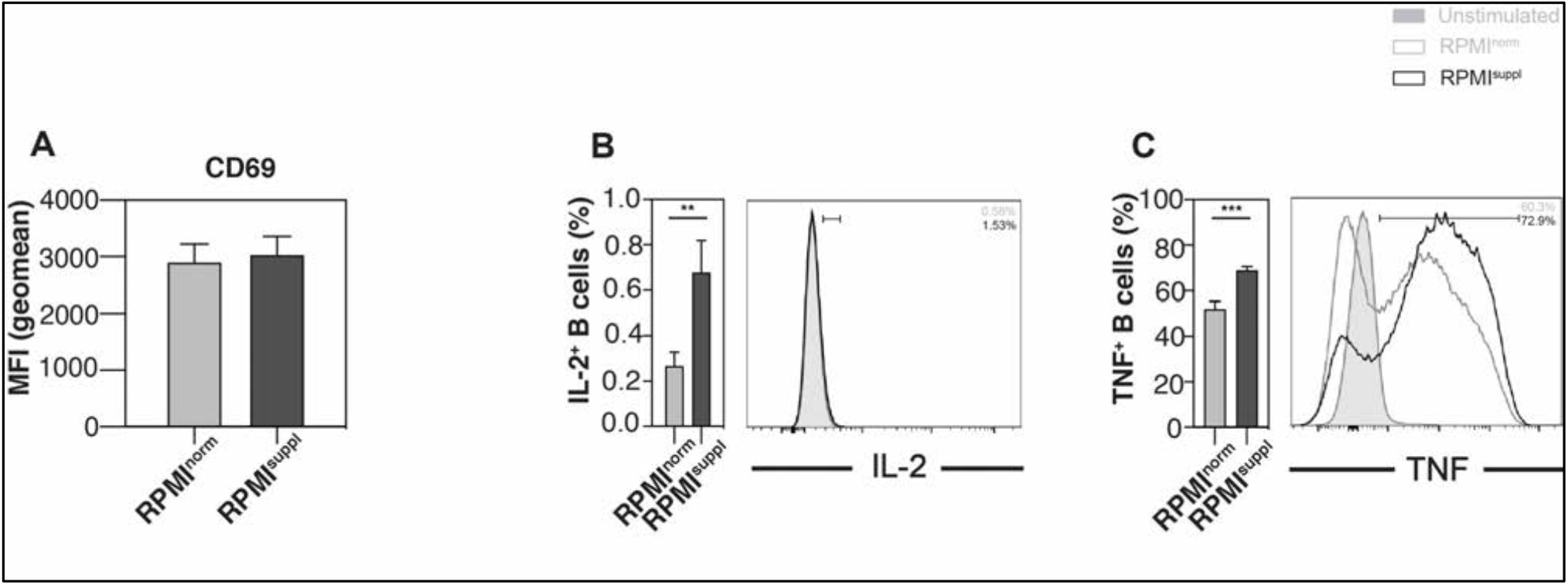
Ca^2+^ supplementation *in vitro* increases the cytokine production of primary human B cells. PBMCs were isolated from residual leukocyte units of healthy donors. PBMCs were stimulated for 4 h with 25 ng/ml PMA and 1 μg/ml ionomycin in either normal RPMI1640 medium (RPMI^norm^) or RPMI1640 medium supplemented with 1 mM Ca^2+^ (RPMI^suppl^). Human B cells (CD14^-^ CD3^-^ CD20^+^ cells) were analysed for the expression of **(A)**CD69 and the production of **(B)** IL-2, and **(C)** TNF. Summary graphs (left panels) and representative data (right panels) from gated B cells are shown, respectively. Data were pooled from four independent experiments with three samples each (n = 12).

**Figure 8:**
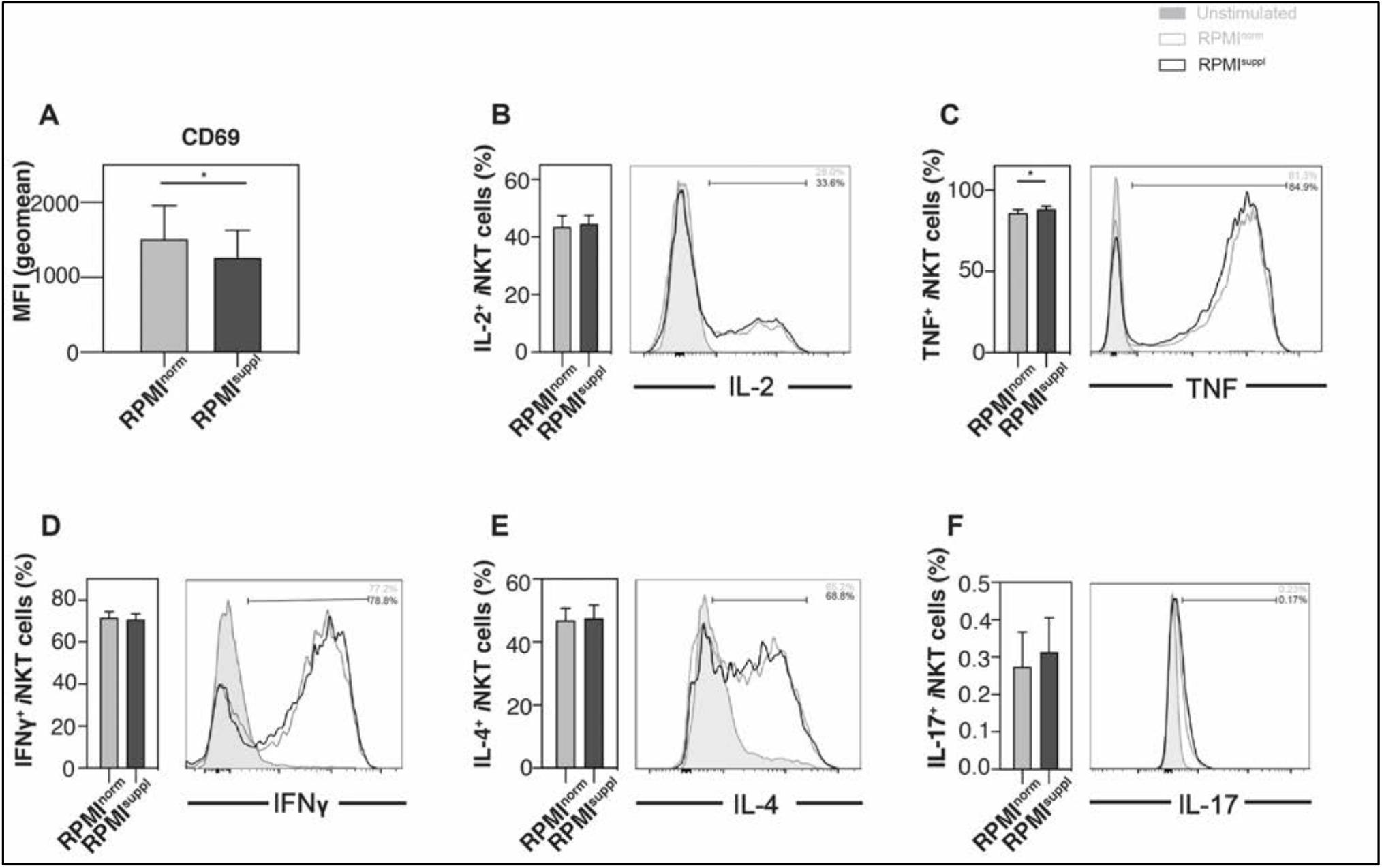
Ca^2+^ supplementation *in vitro* increases TNF production by expanded human *i*NKT cells. *i*NKT cells were expanded *ex vivo* in the presence of αGalCer. The expanded cells were stimulated for 4 h with 25 ng/ml PMA and 1 μg/ml ionomycin in either normal RPMI1640 medium (RPMI^norm^) or RPMI1640 medium supplemented with 1 mM Ca^2+^ (RPMI^suppl^). Vα24*i* NKT cells (live CD14^-^ CD20^-^ CD3^+^ 6B11^+^ cells) were analysed for the expression of (**A**) CD69 and the production of (**B**) IL-2, (**C**) TNF, (**D**) IFNγ, (**E**) IL-4, and **(F)** IL-17 were measured by ICCS. Summary graphs (left panels) and representative data (right panels) from gated *i*NKT cells are shown, respectively. Data were pooled from four independent experiments with three samples each (n = 12).

**Figure 9:**
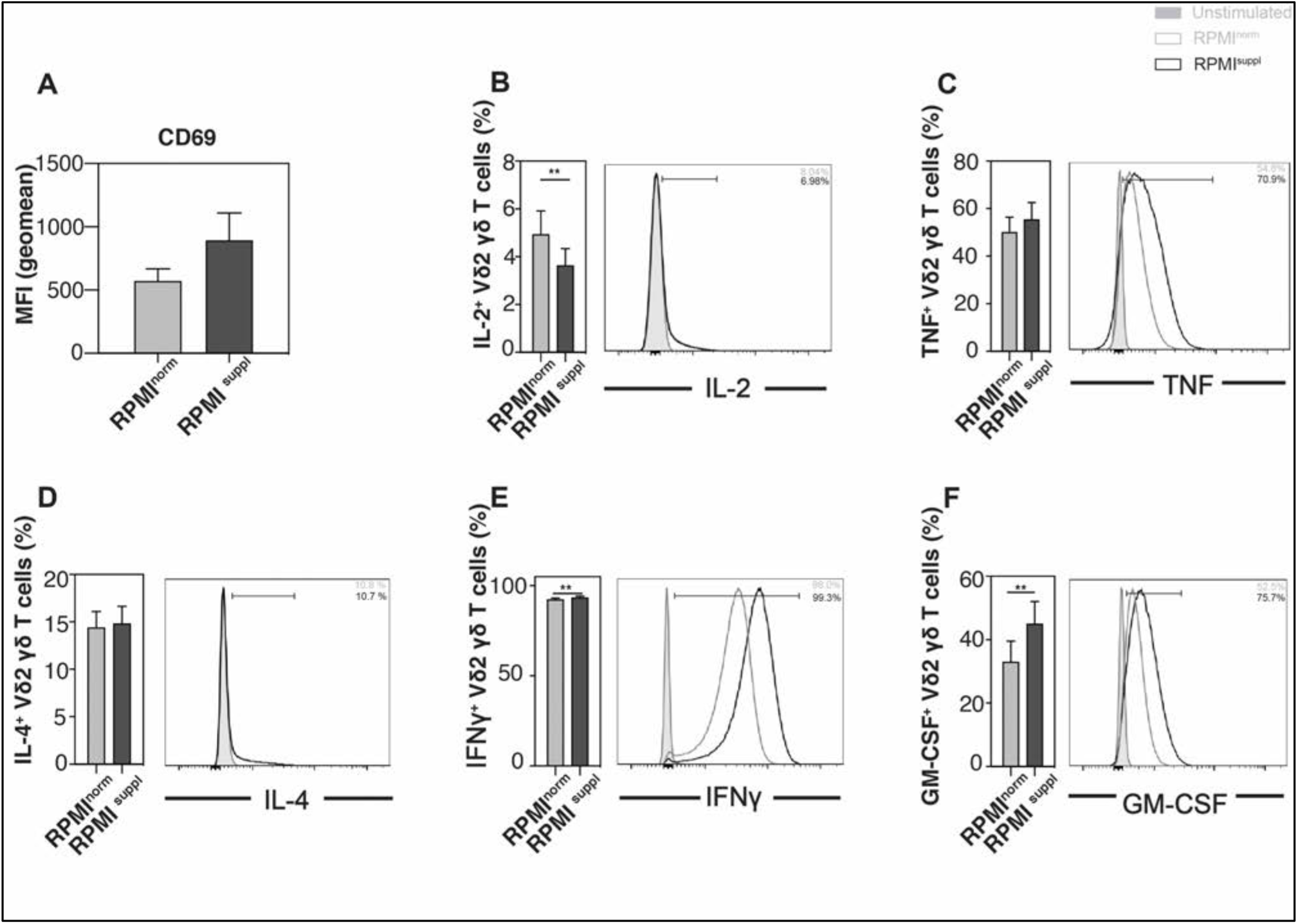
Ca^2+^ supplementation *in vitro* increases the production of some cytokines by expanded human Vγ2^+^ T cells. Vγ2^+^ T cells were expanded *in vitro* in the presence of Zoledronic acid. The expanded cells were stimulated for 4 h with 25 ng/ml PMA and 1 μg/ml ionomycin in either normal RPMI1640 medium (RPMI^norm^) or RPMI1640 medium supplemented with 1 mM Ca^2+^ (RPMI^suppl^). Human Vδ2^+^ T cells (live CD14^-^ CD20^-^ CD3^+^ Vγ2^+^ cells) were analysed for the expression of **(A)** CD69 and the production of **(B)** IL-2, **(C)** TNF, **(D)** IL-4, **(E)** IFNγ, and **(F)** GM-CSF. Summary graphs (left panels) and representative data (right panels) from gated Vγ2^+^ T cells are shown, respectively. Data were pooled from three independent experiments with three samples each (n = 9).

### Increased Ca^2+^ during *in vitro* stimulation of macrophages increases cell death and induces a shift toward M1 polarization

Given the clear ability of increased *in vitro* Ca^2+^ concentrations to augment the cytokine production of stimulated mouse and human lymphoid cells, we tested next the impact on myeloid cells. To this end, we initially measured the impact of elevated Ca^2+^ levels on primary human PBMC monocytes and noticed that some (IFNγ, TNF) but not all (IL-2) cytokines were boosted by the Ca^2+^ supplementation (**Figure 10A-D**), without impact on cell survival (**Supplementary figure 2E**). These data suggested that the cytokine detection by primary myeloid cells could benefit from *in vitro* Ca^2+^ supplementation as well. Blood monocytes have the ability to differentiate into functionally distinct macrophages subsets and, therefore, we tested next the impact of the Ca^2+^ concentration on human monocyte-derived macrophages. Primary human monocyte-derived macrophages (M0 macrophages) were polarized into M1 with LPS and IFNγ, into M2a with IL-4 stimulation, or into M2c with IL-10 with or without Ca^2+^ supplementation. Surprisingly, and in contrast to the findings with human lymphoid cells (**Supplementary figure 2**), the polarization of M0 macrophages in elevated Ca^2+^ concentrations increased cell death, regardless of the subtype they were polarized into (**Figure 11A-C**). Similar results were seen when M0 macrophages were cultured alone in Ca^2+^ supplemented medium (**Supplementary Figure 3**). When M0 macrophages were cultured alone or when they were polarized into M1 macrophages, higher Ca^2+^ levels increased the expression of the M1 markers HLA-DRα and CD86 (**Supplementary Figure 4A**). In contrast, when M0 macrophages were cultured alone or when they were polarized into M2a macrophages, Ca^2+^ supplementation decreased the expression of the M2a markers CD200R and CD206 (**Supplementary Figure 4B**). Importantly, when the expression of the M1 markers HLA-DRα and CD86 was analysed on M2a and M2c macrophages, Ca^2+^ supplementation increased the expression of HLA-DRα on M2a macrophages (**Figure 12A**) and of CD86 on both M2 macrophages subsets (**Figure 12B, C**). These data indicate that increased Ca^2+^ concentrations support phenotypic M1 polarization of human monocyte-derived macrophages *in vitro*. However, these phenotypic changes appeared not to translate into functional changes, as the production of TNF and CXCL10 by M1 macrophages (**Figure 13A, B**) and the production of TGFβ and IL-4 by M2a and M2c macrophages (**Figure 13C, D**) was not influenced by the Ca^2+^ supplementation. These data suggest that for myeloid cells the impact of an increased Ca^2+^ concentration on the cytokine production depends on the activation status of the cells and needs to be tested cell type specifically.

**Figure 10:**
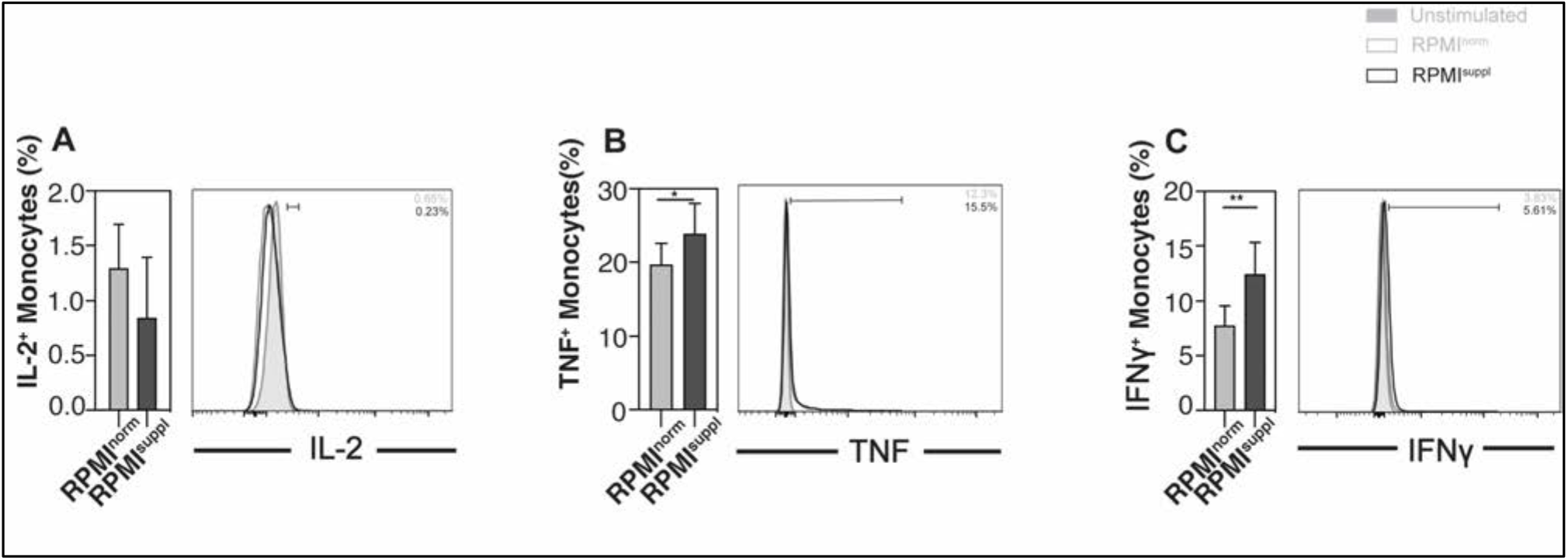
Ca^2+^ supplementation *in vitro* increases the cytokine production of primary monocytes. PBMCs were isolated from residual leukocyte units of healthy donors. PBMCs were stimulated for 4 h with 25 ng/ml PMA and 1 μg/ml ionomycin in either normal RPMI1640 medium (RPMI^norm^) or RPMI1640 medium supplemented with 1 mM Ca^2+^ (RPMI^suppl^). Human monocytes (CD3^-^ CD20^-^ CD14^+^ cells) were analysed for the expression of **(A)** CD69 and the production of **(B)** IL-2, **(C)** TNF, and **(D)** IFNγ. Summary graphs (left panels) and representative data (right panels) from gated monocytes are shown, respectively. Data were pooled from four independent experiments with three samples each (n = 12).

**Figure 11:**
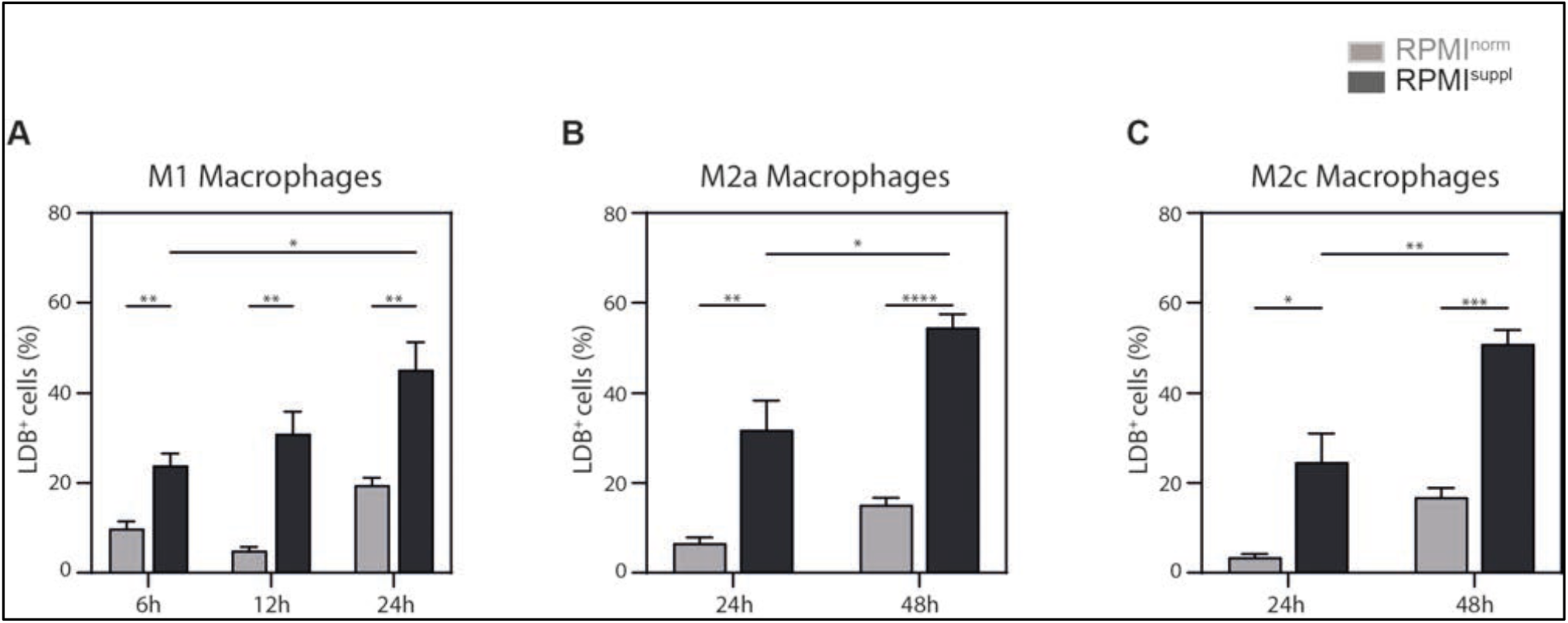
Ca^2+^ supplementation *in vitro* increases cell death in M1, M2a, and M2c polarized macrophages. Primary human monocyte-derived macrophages were polarized into **(A)** M1 (100ng/ml LPS, 20 ng/ml IFNγ), **(B)** M2a (20ng/ml IL-4), or **(C)** M2c (20 ng/ml IL-10) macrophages for the indicated hours in either normal RPMI1640 medium (RPMI^norm^) or RPMI1640 medium supplemented with 1 mM Ca^2+^ (RPMI^suppl^). The bar graphs show the relative percentages of cells positive for LIVE/DEAD Fixable Blue Dead Cell Stain (LDB^+^ cells), indicating dead cells. The data shown are means ± SEM of biological replicates of six donors, pooled from two independent experiments with similar results.

**Figure 12:**
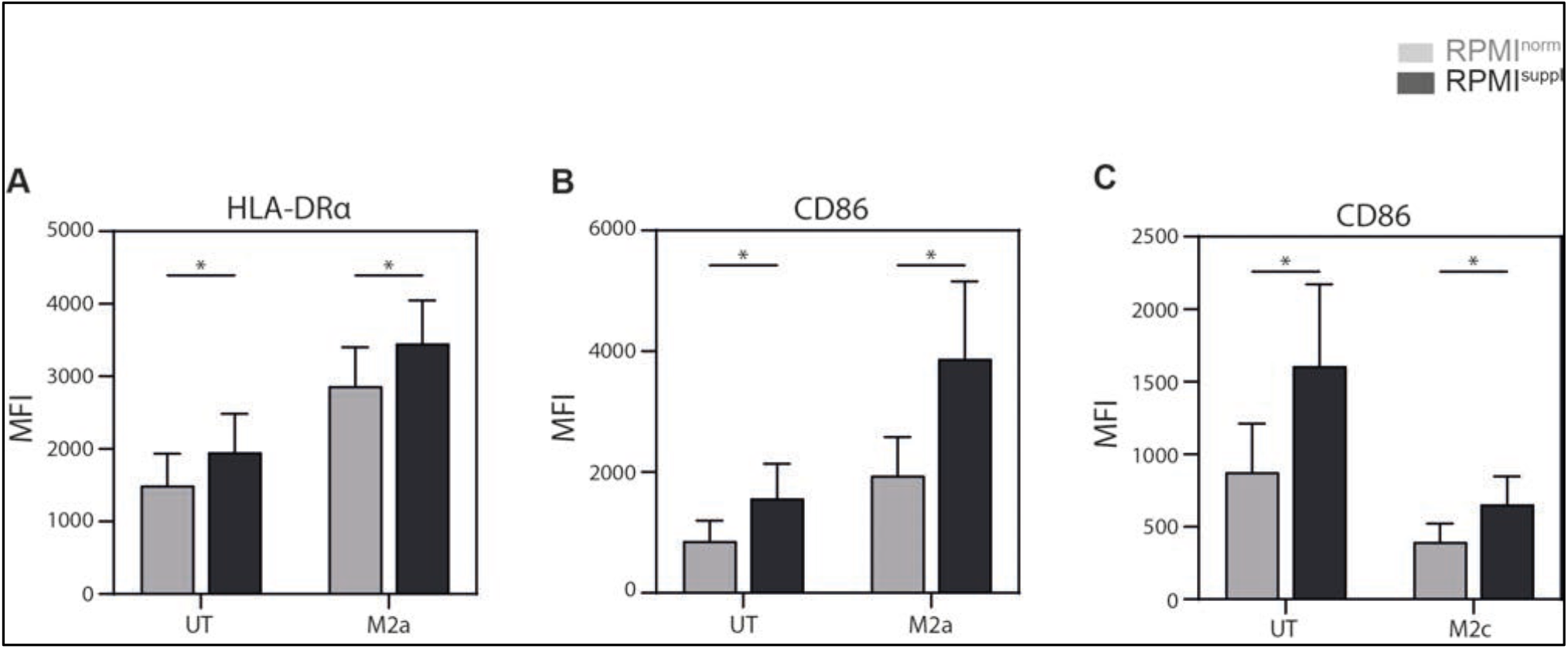
Ca^2+^ supplementation *in vitro* increases the expression of cell surface M1 markers on M2 macrophages. Primary human monocyte-derived macrophages were cultured untreated (UT) or polarized into (A, B) M2a (20 ng/ml IL-4, 24 hours) or into (C) M2c (20ng/ml IL-10, 24 hours) macrophages in either normal RPMI1640 medium (RPMI^norm^) or RPMI1640 medium supplemented with 1 mM Ca^2+^ (RPMI^suppl^). The expression (mean fluorescent intensity, MFI) of **(A)** HLA-DRα (UT, M2a) and **(B, C)** CD86 ((B): UT, M2a; (C): UT, M2c) on live macrophages is shown. The data shown are means ± SEM of biological replicates of six donors, pooled from two independent experiments with similar results.

**Figure 13:**
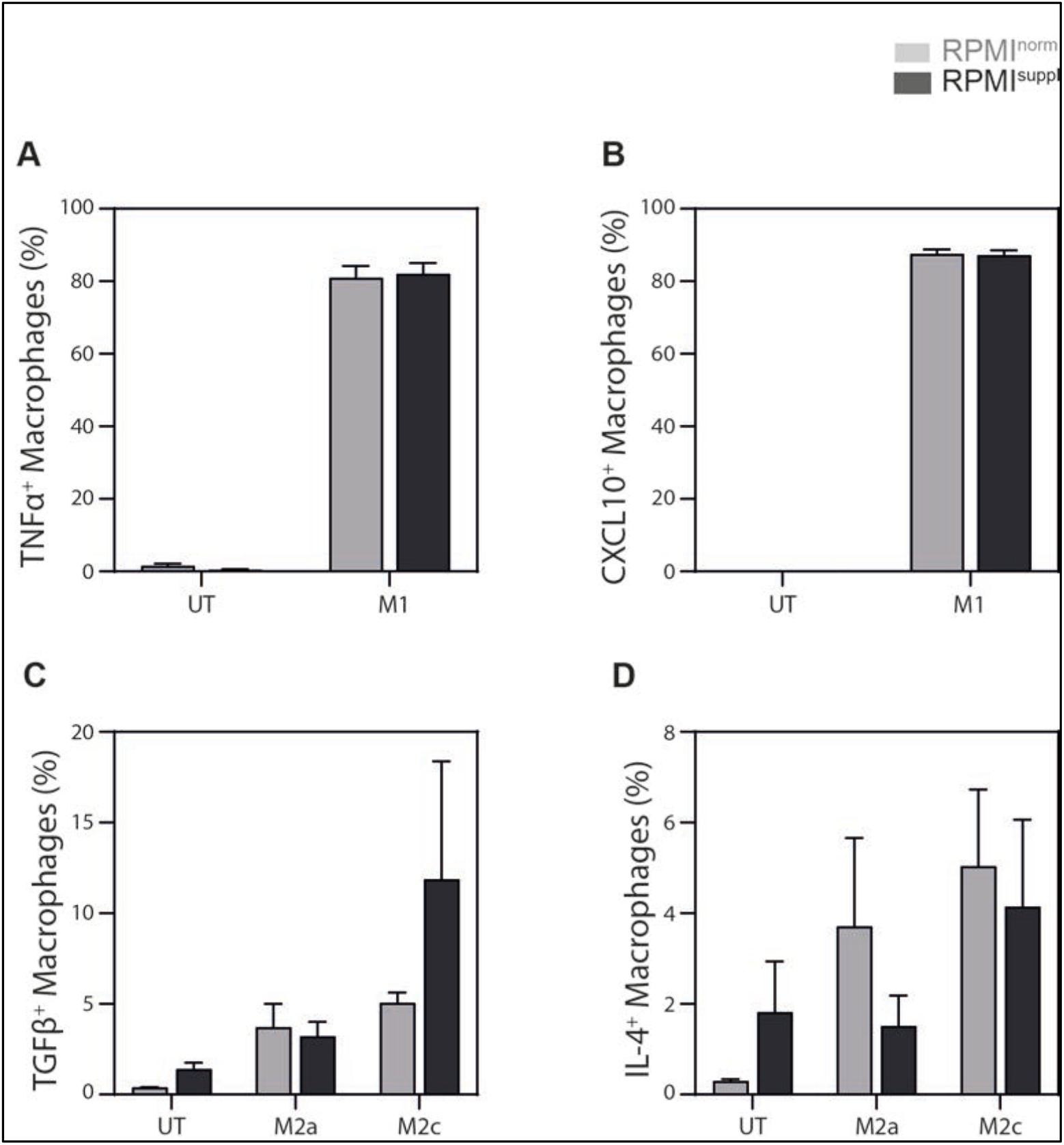
Ca^2+^ supplementation *in vitro* does not impact the cytokines production of macrophages. Primary human monocyte-derived macrophages were cultured untreated (UT) or polarized into M1 (100ng/ml LPS, 20 ng/ml IFNγ, 6 hours), M2a (20 ng/ml IL-4, 48 hours), or M2c (20 ng/ml IL-10, 48 hours) macrophages in either normal RPMI1640 medium (RPMI^norm^) or RPMI1640 medium supplemented with 1 mM Ca^2+^ (RPMI^suppl^). The production of **(A)** TNF, **(B)** CXCL10, **(C)** TGFβ, and **(D)** IL4 by indicated macrophages was analysed ICCS. The data shown are means ± SEM of biological replicates of five (TGFβ, IL-4) or six donors (CXCL10, TNF), pooled from two independent experiments with similar results.

## Discussion

Here we demonstrate that Ca^2+^ supplementation of the RPMI1640 medium to a final concentration of approx. 1.8 mM Ca^2+^ can increase the cytokine production by both mice and human lymphoid cells during PMA/ionomycin stimulation *in vitro* without impacting the survival of the cells. This effect appeared stronger for primary lymphoid cells than *in vitro* expanded cells. Primary human monocytes responded similarly with an augmented cytokine production to the elevated Ca^2+^ concentrations. However, for human monocyte-derived macrophages a distinct effect was observed: elevated Ca^2+^ concentrations *in vitro* led to increased cell death and promoted phenotypic M1 polarization without impacting cytokine production.

Previous work on conventional mouse^4^ and human^5^ αβ T cells showed that the 0.49 mM Ca^2+^ in RPMI1640 are suboptimal to drive cytokine production during *in vitro* stimulation and a minimal concentration of 1.5 mM was suggested. The basal level of intracellular Ca^2+^ in T cells is approx. 100 nM and can increases to 1 μM following stimulation^22^. Therefore, it is not self-evident how an increase of the extracellular Ca^2+^ concentration from 0.49 mM to approx. 1.8 mM can boost cytokine production by conventional αβ T cells^4,5^. Irrespective of the mechanism, this observation is important to optimize the analysis of conventional αβ T cells *in vitro*. Here, we confirmed that unconventional αβ T cells, γδ T cells, and B cells from both mouse and human also produce more cytokines after *in vitro* stimulation in the presence of elevated Ca^2+^ levels. Considering the Ca^2+^ found in FCS and human serum^5^, we estimated that it added around 0.3 mM Ca^2+^ to our medium. Therefore, the complete RPMI (RPMI^norm^) and the Ca^2+^ supplemented (RPMI^suppl^) medium contained approx. 0.7 - 0.8 mM and 1.7 - 1.8 mM Ca^2+^, respectively. The Ca^2+^ concentration of the extracellular fluid *in vivo* in humans was measured to be 2.2 - 2.7 mM^21^. This indicates that the Ca^2+^ supplementation brings the Ca^2+^ concentration of the RPMI^suppl^ medium close to the physiological conditions *in vivo*.

For primary mouse lymphoid cells, the elevated Ca^2+^ concentration increased the production of 6 out of 6 cytokines tested for *i*NKT cells (Figure 1), 3 out of 5 for γδ T cells (Figure 2), and 2 out of 3 for B cells (Figure 3), without a decrease of the production of any of the cytokines tested. Interestingly, the boosting Ca^2+^ effect appeared weaker for primary human lymphoid cells: the elevated Ca^2+^ concentration increased the production of 1 out of 5 cytokines tested for *i*NKT cells (Figure 4), 2 out of 4 for γδ T cells (Figure 5), 3 out of 4 for MAIT cells (Figure 6), and 2 out of 2 for B cells (Figure 7), again without any decreases of any of the cytokines tested. The reason for this species difference is unclear at this point.

The cytokine response of the *in vitro* expanded cell lines under increased Ca^2+^ concentrations was comparable to the primary cells: both primary and expanded *i*NKT cells showed increased production for 1 (TNF) out of 5 cytokines tested (Figure 4 and 8). Primary γδ T cells showed increased production for 2 (IFNγ, TNF) out of 4 (Figure 5) and expanded for 2 (GM-CSF, IFNγ) out of 5 (Figure 9) cytokines tested. Surprisingly, expanded γδ T cells showed a decrease of IL-2 with increased Ca^2+^ (Figure 9), which is the only instance in which we noticed a decrease in cytokine production.

Similar to the lymphoid cells, primary human monocytes increased the production of 2 out of 3 cytokines tested (Figure 10) when stimulated in the presence of elevated Ca^2+^ levels (Figure 10). However, the data we obtained with human primary monocyte-derived macrophages were in contrast to the ones from lymphoid cells and primary human monocytes. Most importantly, we noticed a clear increase in the frequency of macrophages that died when incubated for more than six hours in the presence of elevated Ca^2+^ levels (Figure 11, Supplementary Figure 3). Calcium-induced ER-stress^23,24^ and mitochondrial changes^25^ can trigger ROS-(reactive oxygen species) production in macrophages, which can lead to cell death^23,24,26–28^. This might explain why macrophages are more sensitive to the Ca^2+^ concentration of the medium. Furthermore, we observed a shift towards an M1 phenotype for both M0 macrophages (Supplementary Figure 4) as well as M1 and M2 macrophages (Figure 12, Supplementary Figure 4) in the presence of elevated Ca^2+^ levels. However, these phenotypic changes appeared not to translate into functional changes (Figure 13). Incidentally, ROS-induced NFκB/MAPK activation in macrophages^29,30^ supports M1 polarization^31–33^. However, given the large amount of macrophage cell death we observed, we cannot exclude the possibility that the apparent M1 shift is the result of M1 macrophages potentially being less sensitive to this Ca^2+^-induced cell death. We are aware of only two other studies on the impact of Ca^2+^ on the cytokine production of macrophages. Both showed that calcium influx can impair LPS-induced IL-12 production of mouse macrophages^34,35^ without affecting the production of TNF or IL-6^35^. Therefore, it is unclear at this stage whether elevated Ca^2+^ concentration *in vitro* can influence the cytokine production of macrophages.

In summary, our data demonstrate that the Ca^2+^ concentration during *in vitro* cultures is an important variable to be considered for functional experiments and that a concentration of approx. 1.8 mM Ca^2+^ can boost the cytokine production by both mice and human lymphoid cells.

## Material and Methods

### Human samples

Residual leukocyte units from healthy donors were obtained from the Dokuz Eylul University Blood Bank (Izmir, Turkey) after written consent. The ethical approval for the study was obtained from the ‘Noninvasive Research Ethics Committee’ of the Dokuz Eylul University (approval number: 2018/06-27/3801-GOA).

### Mice

All mice were housed in the vivarium of the Izmir Biomedicine and Genome Center (IBG, Izmir, Turkey) in accordance with the respective institutional animal care committee guidelines. C57BL/6 and BALB/c mice were originally purchased from the Jackson Laboratories (Bar Harbor, USA). All mouse experiments were performed with prior approval by the institutional ethic committee (‘Ethical Committee on Animal Experimentation’), in accordance with national laws and policies. All the methods were carried out in accordance with the approved guidelines and regulations.

### Reagents, monoclonal antibodies, and flow cytometry

α-galactosylceramide (αGalCer) was obtained from Avanti Polar Lipids (USA). Monoclonal antibodies against the following mouse antigens were used in this study: CD3ε (145-2C11), CD8α (53-6.7), CD4 (RM4-5), CD44 (IM7), CD45R (RA3-6B2), CD90.2 (30-H12), CD117 (2B8), CD127 (A7R34), γδTCR (UC7-13D5), GM-CSF (MP1-22E9), IFNγ (XMG1.2), IL-2 (JES6-5H4), IL-4 (11B11), IL-10 (JES5-16E3), IL-13 (eBio13A), IL-17A (TC11-18H10), Ly6A/E (D7), Ly6G (1A8), and NK1.1 (PK136). Monoclonal antibodies against the following human antigens were used in this study: CD3 (UCHT1), CD14 (M5E2), CD20 (2H7), CD64 (10.1), CD69 (FN50), CD86 (IT2.2), CD161 (HP-3G10), CD163 (GHI/61), CD200R (OX-108), CD206 (15-2), CXCL10 (33036), HLA-DR (L243), IFNγ (4S.B3), IL-2 (MQ1-17H12), IL-4 (8D4-8), IL-10 (JES3-9D7), IL-17 (64DEC17), *i*Vδ24 (6B11), pan-γδTCR (B1), TGFβ (TW4-2F8), TNF (Mab11), Vδ2 TCR (B6), and Vα7.2 (3C10), Antibodies were purchased from either BioLegend (USA), BD Biosciences (USA), or eBiosciences (USA). Antibodies were conjugated to Brilliant Ultra Violet 395, Pacific Blue, Violet 500, Brilliant Violet 570, Brilliant Violet 605, Brilliant Violet 650, Brilliant Violet 711, Brilliant Violet 785, FITC, PerCP-Cy5.5, PerCP-eF710, PE, PE-CF594, PE-Dazzle594, PE-Cy7, APC, AF647, eF660, AF700, APC-Cy7, or APC-eF780. Anti-mouse CD16/32 (2.4G2) antibody (Tonbo Biosciences, USA) or Human TruStain FcX (BioLegend) was used to block Fc receptors. Unconjugated mouse and rat IgG antibodies were purchased from Jackson ImmunoResearch (USA). Dead cells were labelled with Zombie UV Dead Cell Staining kit (BioLegend) or with LIVE/DEAD Fixable Blue Dead Cell Stain kit (ThermoFisher, USA). Flow cytometry and preparation of fluorochrome-conjugated antigen-loaded CD1d tetramers were performed as described^36^. Graphs derived from digital data are displayed using a ‘bi-exponential display’. Cell were gated as follows: (a) mouse: Vα14*i* NKT cells (live CD8α^-^CD19/CD45R^-^ CD44^+^ TCRβ/CD3ε^+^ CD1d/PBS57-tetramer^+^ cells), γδ T cells (live CD19/CD45R^-^ CD4^-^ CD8α^-^CD3ε^+^ γδTCR^+^ cells), and B cells (live CD3ε^-^CD4^-^ CD8α^-^CD19/CD45R^+^ cells); (b) human: Vα24*i* NKT cells (live CD14^-^ CD20^-^ CD3^+^ 6B11^+^), δ2^+^ T cells *ex vivo* (live CD14^-^ CD20^-^ CD3^high^ γδTCR^low^ cells, or live CD14^-^ CD20^-^ CD3^+^ Vδ2^+^ cells) and *in vitro* (live CD14^-^ CD20^-^ CD3^+^ Vδ2^+^ cells), MAIT cells (live CD14^-^ CD20^-^ CD3^+^ Vα7.2^+^ CD161^+^ cells), B cells (live CD3^-^ CD14^-^ CD20^+^ cells), and macrophages (live CD68^+^ cells).

### Cell preparation

Single-cell suspensions from mouse spleens were prepared as described^37^. Red blood cells and dead cells were eliminated through Lymphoprep (StemCell Technologies, Canada) density gradient centrifugation, as previously described^37^. Human PMBCs were obtained from the residual leukocyte units of healthy donors via density gradient centrifugation with Ficoll-Paque Plus (GE Healthcare, USA). To obtain monocytes, a second density gradient centrifugation with Percoll (GE Healthcare) was performed.^38^

### *In vitro* expansion of *i*NKT cells

Human *i*NKT cells were expanded from resting PBMCs of healthy donors as described before^39^. Briefly, freshly isolated PBMCs (1 × 10^6^ cell/ml, 5 ml/well) were treated with 100 ng/ml αGalCer (KRN7000, Avanti Polar Lipids) and cultured for 13 days. 20 IU/ml human recombinant IL-2 (Proleukin, Novartis) was added to the cultures every other day starting from day 2. From day 6 onwards, the concentration of IL-2 was increased to 40 IU/ml. At the end of the expansion, an aliquot of each sample was collected and analysed for *i*NKT cell expansion by flow cytometry. Expansion was done in RPMI 1640 (Gibco or Lonza) supplemented with 5% (v/v) Human AB serum (Sigma-Aldrich), 1% (v/v)

Penicillin/Streptomycin, 1 mM sodium pyruvate (Lonza), 1% (v/v) non-essential amino acids (Cegrogen), 15 mM HEPES buffer (Sigma-Aldrich), and 55 μM 2-mercaptoethanol (AppliChem).

### *In vitro* expansion of Vδ2^+^ T cells

Human Vδ2^+^ T cells were expanded from PBMCs of healthy donors similar to published protocols^39–41^. Briefly, freshly isolated PBMCs (1 × 10^6^ cell/ml, 5 ml/well) were cultured with 5 μM Zoledronic acid (Zometa, Novartis) in the presence of 100 IU/ml human recombinant IL-2 (Proleukin, Novartis) for 13 days. IL-2 was replenished every other day and from day 6 onwards the concentration was increased to 200 IU/ml. The cultures were performed in RPMI1640 (Gibco or Lonza) supplemented with 5% (v/v) Human AB serum (Sigma-Aldrich), 1% (v/v) Penicillin/Streptomycin (Gibco), 1 mM sodium pyruvate (Lonza), 1% (v/v) non-essential amino acids (Cegrogen), 15 mM HEPES buffer (Sigma-Aldrich), and 55 μM 2-mercaptoethanol (AppliChem).

### Macrophage generation

Purified monocytes were cultured in RPMI1640 (Gibco or Lonza) medium containing 5% FBS (Mediatech), 1% Penicillin/Streptomycin (Gibco), and 10 ng/ml human recombinant M-CSF (Peprotech) for their differentiation into macrophages. 3 × 10^6^ cells/well were seeded in low-attachment 6-well plates (Corning Costar) and incubated for 7 days at 5% CO_2_ and 37°C. Macrophages were collected and cultured in 24-well cell culture plates as 5 × 10^5^ macrophages/well for 24 hours in RPMI1640 medium containing 5% FBS, 1% Penicillin/Streptomycin. On the following day, the cell culture media was replaced with fresh media and macrophages were stimulated with the relevant polarization factors as described below. The cells were verified to be over 90% CD68^+^ by flow cytometry.

### *In vitro* stimulation

Splenocytes were stimulated *in vitro* with 50 ng/ml PMA and 1 μg/ml ionomycin (both Sigma-Aldrich) for 4 h at 37°C in the presence of both Brefeldin A (GolgiPlug) and Monensin (GolgiStop, both BD Biosciences). As GolgiPlug and GolgiStop were used together, half the amount recommended by the manufacturer where used, as suggested previously^42^. Cells were stimulated in complete RPMI medium (RPMI 1640 supplemented with 10% (v/v) FCS (Mediatech), 1% (v/v) Pen-Strep-Glutamine (10.000 U/ml penicillin, 10.000 μg/ml streptomycin, 29.2 mg/ml L-glutamine (Invitrogen)) and 50 μM β-mercaptoethanol (AppliChem)). For human studies, freshly isolated PBMCs or expanded cell populations were stimulated *in vitro* with 25 ng/ml PMA and 1 μg/ml PMA for 4 h at 37°C in the presence of Brefeldin A or monensin. Cells were stimulated in complete RPMI medium (RPMI 1640 supplemented with 10% (v/v) FBS, 1% (v/v) Penicillin/Streptomycin, 1 mM sodium pyruvate (Lonza), 1% (v/v) non-essential amino acids (Cegrogen), 15 mM HEPES buffer (Sigma-Aldrich), and 55 μM 2-mercaptoethanol (AppliChem). The complete RPMI medium containing FCS contains approx. 0.7 - 0.8 mM Ca^2+^ (RPMI^norm^). To obtain the RPMI^suppl^ medium containing elevated Ca^2+^ concentrations, 1 mM CaCl_2_ (Sigma-Aldrich) was added to the complete RPMI medium.

### Macrophage polarization

Macrophages were stimulated (i) for M1 polarization with 100 ng/ml LPS (InvivoGen) and 20 ng/ml IFNγ (R&D); (ii) for M2a with 20 ng/ml IL-4 (R&D); or (iii) for M2c with 20 ng/ml IL-10 (R&D). The incubation times are specified in the text.

### Statistical analysis

Results are expressed as mean ± standard error of the mean (SEM). Statistical comparisons were drawn using a two-tailed paired Student t-test (Excel, Microsoft Corporation, USA; GraphPad Prism, GraphPad Software, USA) for all normally distributed paired samples, a Wilcoxon matched-pairs signed rank test (GraphPad Prism) for non-normally distributed samples or otherwise using an ANOVA test (GraphPad Prism). Normal distribution was tested using D’Agastino – Pearson Test. Statistical analyses were performed with either two-tailed paired students’ T test (normally distributed data) or Wilcoxon matched-pairs signed rank test (not normally distributed data). p-values <0.05 were considered significant and are indicated with (*p < 0.05, **p < 0.01, ***p < 0.001, ****p <0.0001. Each experiment was repeated at least twice and background values were subtracted. Graphs were generated with GraphPad Prism (GraphPad Software).

## Abbreviations

αGalCer: α-galactosylceramide
FCS: fetal calf serum
ICCS: intracellular cytokine staining
*i*NKT: invariant Natural Killer T
MAIT: mucosal-associated invariant T
MFI: mean fluorescent intensity
PBMCs: peripheral blood mononucleated cells
ROS: reactive oxygen species.

## Acknowledgments

The authors wish to thank the Flow Cytometry Core Facility and the vivarium at the Izmir Biomedicine and Genome Center (IBG) for excellent technical assistance. We are grateful to the NIH Tetramer Core Facility for providing the mouse CD1d/PBS57 tetramers. This work was funded by grants from TUBITAK (117Z216, GW) and the European Molecular Biology Organization (EMBO, Installation Grant #3073, GW). The funders had no role in study design, data collection and analysis, decision to publish, or preparation of the manuscript.

## Author contributions

Conceived and designed the experiments: YCE, ZOA, SG, AK, DS, GW. Performed the experiments: YCE, ZOA, SG, AK, DGH. Analysed the data: YCE, ZOA, SG, AK, DS, GW. Provided essential reagents: YD. Wrote the paper: YCE, ZOA, DS, GW. All authors reviewed and approved the manuscript.

## Additional information

Supplementary information accompanies this paper at doi:

Competing financial interests: The authors declare no competing financial interests.

## Figures and legends

**Supplementary figure 1:**
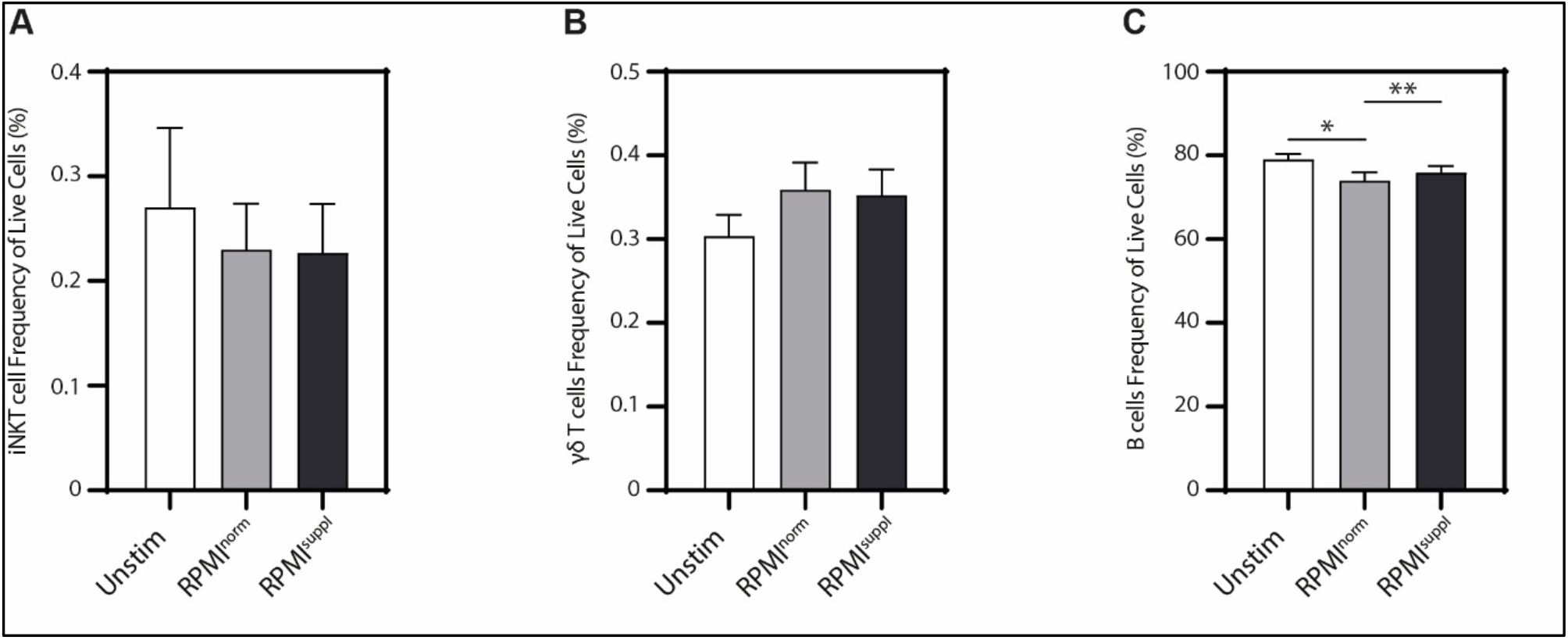
Stimulation of murine *i*NKT cells, γδ T cells, and B cells in RPMI^suppl^ has no negative impact on cell viability. Splenocytes from C57BL/6 mice were stimulated 4 h with 50 ng/ml PMA and 1 μg/ml ionomycin in either normal RPMI1640 medium (RPMI^norm^) or RPMI1640 medium supplemented with 1 mM Ca^2+^ (RPMI^suppl^). **(A)** *i*NKT cells; **(B)** Vδ2^+^ T cells; and **(C)** B cells were stained and analysed by flow cytometry. The bar graphs show the relative percentages of cells positive for LIVE/DEAD Fixable Blue Dead Cell Stain, indicating dead cells. Data were pooled from three independent experiments with three mice per group per experiment (n = 9).

**Supplementary figure 2:**
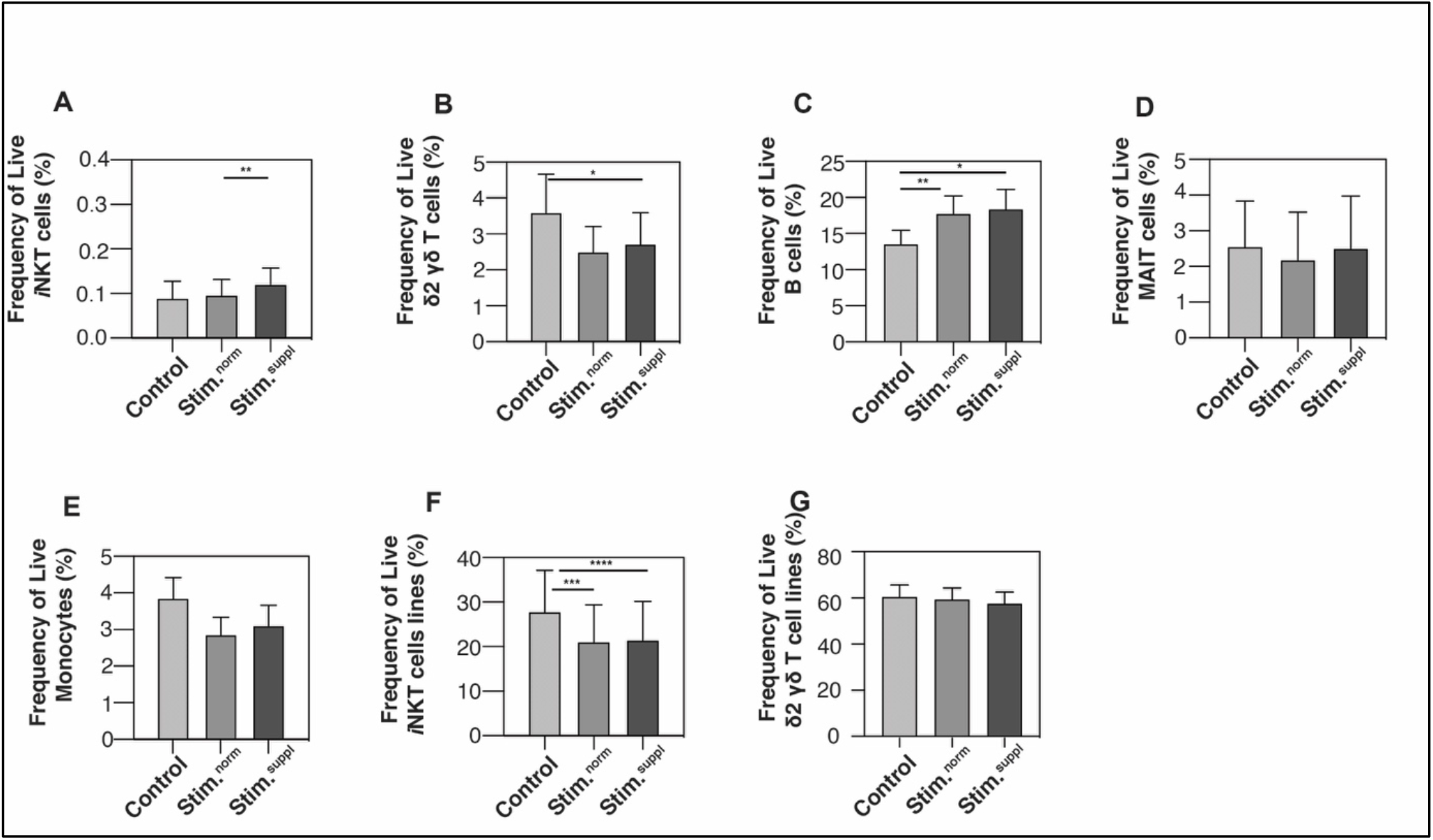
Stimulation of human lymphoid cells in RPMI^suppl^ has no negative effect on cell viability. PBMCs were isolated from residual leukocyte units of healthy donors and were stimulated either directly (A-D) or after in vitro expansion of indicated cells (E, F). The cells were stimulated for 4 h with 25 ng/ml PMA and 1 μg/ml ionomycin in either normal RPMI1640 medium (RPMI^norm^) or RPMI1640 medium supplemented with 1 mM Ca^2+^ (RPMI^suppl^). Primary **(A)** *i*NKT cells, **(B)**Vδ2^+^ T cells, **(C)** B cells; and **(D)** MAIT cells, **(E)** Monocytes or *in vitro* expanded **(F)** *i*NKT cells and **(G)** Vδ2^+^ T cells were stained and analysed by flow cytometry. The bar graphs show the relative percentages of cells positive for LIVE/DEAD Fixable Blue Dead Cell Stain, indicating dead cells. Data were pooled from three (A-B and G) or four (C to F) independent experiment with three samples each (n = 9 - 12).

**Supplementary figure 3:**
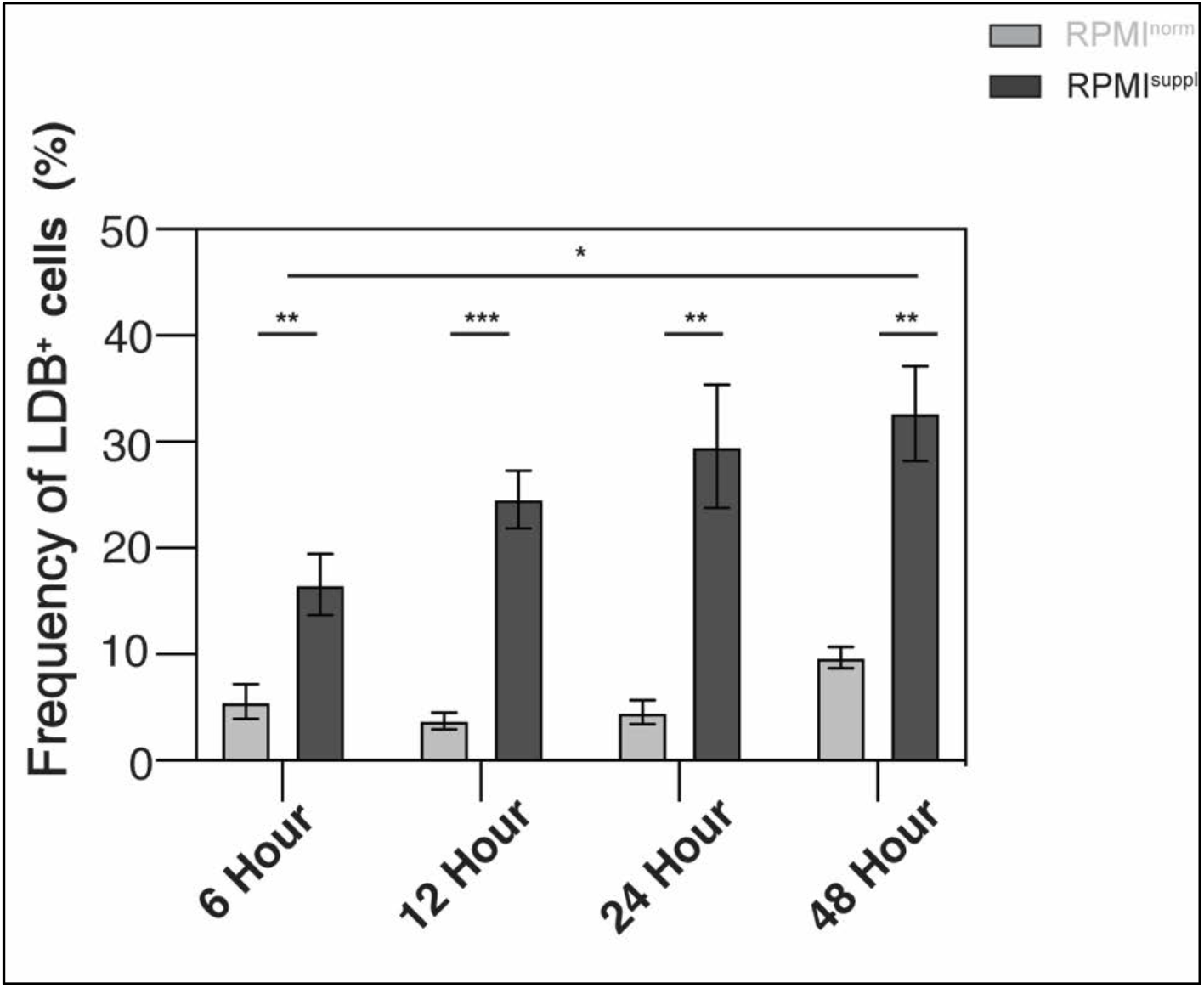
Ca^2+^ supplementation *in vitro* increases cell death of M0 macrophages. Primary human monocyte-derived macrophages were cultured for 6, 12, 24, or 48 hours in either normal RPMI1640 medium (RPMI^norm^) or RPMI1640 medium supplemented with 1 mM Ca^2+^ (RPMI^suppl^). The bar graphs show the relative percentages of cells positive for LIVE/DEAD Fixable Blue Dead Cell Stain (LDB^+^ cells), indicating dead cells. Data shown are means ± SEM of biological replicates of six donors, pooled from two independent experiments with similar results.

**Supplementary figure 4:**
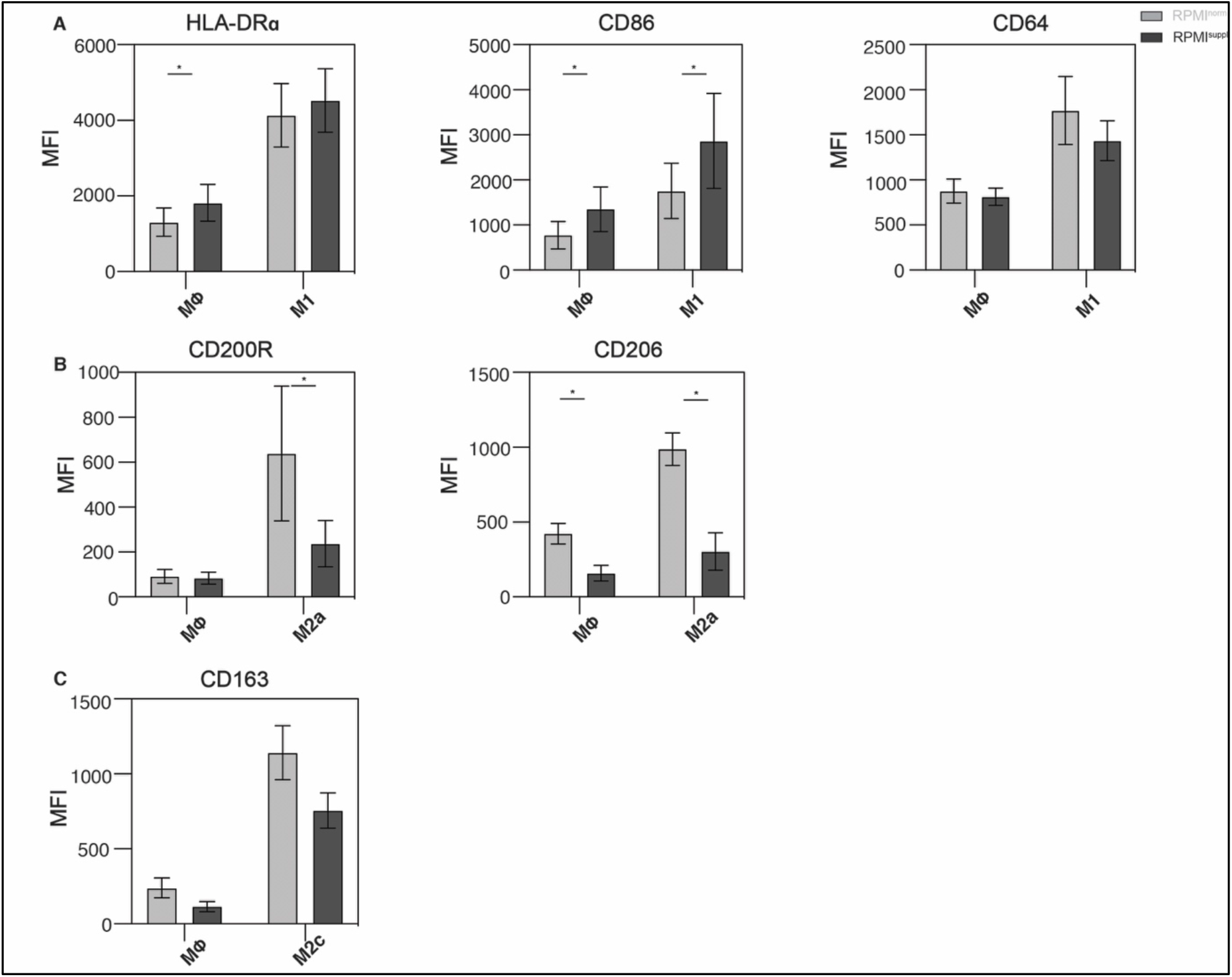
Ca^2+^ supplementation *in vitro* affects the expression of macrophage polarization markers. Primary human monocyte-derived macrophages were cultured untreated (MØ) or polarized into M1 (100ng/ml LPS, 20 ng/ml IFNγ, 12 hours), M2a (20 ng/ml IL-4, 24 hours), or M2c (20ng/ml IL-10, 24 hours) macrophages in either normal RPMI1640 medium (RPMI^norm^) or RPMI1640 medium supplemented with 1 mM Ca^2+^ (RPMI^suppl^). The surface expression (mean fluorescent intensity, MFI) of indicated **(A)** M1 markers (HLA-DR, CD86, CD64), **(B)** M2a markers (CD200R, CD206), and an **(C)** M2c marker (CD163) are shown. The data shown are means ± SEM of biological replicates of six donors, pooled from two independent experiments with similar results.

